# Fine-scale variability in coral bleaching and mortality during a marine heatwave

**DOI:** 10.1101/2022.11.01.514760

**Authors:** Shreya Yadav, Ty NF Roach, Michael J McWilliam, Carlo Caruso, Mariana Rocha de Souza, Catherine Foley, Corinne Allen, Jenna Dilworth, Joel Huckeba, Erika P Santoro, Renee Wold, Jacqueline Simpson, Spencer Miller, Joshua R Hancock, Crawford Drury, Joshua S Madin

## Abstract

Coral bleaching and mortality can show significant spatial and taxonomic heterogeneity at local scales, highlighting the need to understand the fine-scale drivers and impacts of thermal stress. In this study, we used structure-from-motion photogrammetry to track coral bleaching, mortality, and changes in community composition during the 2019 marine heatwave in Kāne‘ohe Bay, Hawai‘i. We surveyed 30 shallow reef patches every 3 weeks for the duration of the bleaching event (August-December) and one year after, resulting in a total of 210 large-area, high-resolution photomosaics that enabled us to follow the fate of thousands of coral colonies through time. We also measured environmental variables such as temperature, sedimentation, depth, and wave velocity at each of these sites, and extracted estimates of habitat complexity (rugosity R and fractal dimension D) from digital elevation models to better understand their effects on patterns of bleaching and mortality. We found that up to 80% of corals experienced moderate to severe bleaching in this period, with peak bleaching occurring in October when heat stress (DHW) reached its maximum. Mortality continued to accumulate as bleaching levels dropped, driving large declines in more heat-susceptible species (77% loss of *Pocillopora* cover) and moderate declines in heat-tolerant species (19% and 23% for *Porites compressa* and *Montipora capitata*, respectively). Declines in live coral were accompanied by a rapid increase in algal cover across the survey sites. Spatial differences in bleaching were significantly linked to habitat complexity and coral species composition, with reefs that were dominated by *Pocillopora* experiencing the most severe bleaching. Mortality was also influenced by species composition, fractal dimension, and site-level differences in thermal stress. Our results show that spatial heterogeneity in the impacts of bleaching are driven by a mix of environmental variation, habitat complexity, and differences in assemblage composition.

## Introduction

Increasingly severe and frequent marine heat waves have caused large-scale losses in coral cover (Heron et al., 2016; Spalding & Brown, 2015). In the past decade alone, consecutive thermal stress events in 2014, 2015, 2016, and 2017 have had global impacts on the structure and functioning of coral reef ecosystems (Arthur et al., 2006; Hughes et al., 2017). Elevated temperatures disrupt the partnership between corals and their algal symbionts (*Symbiodineacea*) causing them to lose their color, and in cases of prolonged stress, die (Hoegh-Guldberg, 1999). Degraded reefs support reduced biodiversity, which has implications for marine food webs, nutrient cycling, and fisheries (Alvarez-Filip et al., 2009; Eddy et al., 2021). Furthermore, they are unable to attenuate wave energy, making coastlines more vulnerable to storms and cyclones (Ferrario et al., 2014). These changes directly affect the social and economic resilience of communities that depend on reef ecosystems for their lives and livelihoods.

Part of the challenge of managing reefs for their resilience is that the effects of elevated temperatures are not homogenous across coral species, habitats, and geographies, with some species and some locations better able to resist and recover from thermal stress than others. Resilience to bleaching involves a range of processes. If corals die, recovery potential is dependent on the replenishment of populations through larval recruitment and growth, which may take decades (Hughes et al., 2000). Yet, corals can also avoid mortality by restoring symbiont populations and regaining colour in months (i.e., individual-level recovery) (Gilmour et al. 2013). Biotic variability in resilience to bleaching is therefore derived from many sources, from regional “supply-side” dynamics to differences in host *Symbiodinium* (Jones et al., 2008) and coral colony size and morphology (Brandt, 2009). Some coral genera like *Acropora* and *Pocillopora* have generally been recorded to be more bleaching susceptible than others but are also often the species that drive recovery after a major disturbance (Burt et al., 2008; Marshall & Baird, 2000). In some cases, corals in deeper reefs have been able to better withstand elevated temperatures than shallower ones (Bridge et al., 2013; Baird et al. 2018) but coral species in some shallow lagoonal environments have also shown a remarkable capacity for resistance to high temperatures through their association with thermally tolerant symbionts and/or via host mediated adaptation and acclimatization (Craig et al., 2001; Cunning et al., 2016; Drury, 2020; Roach et al., 2020).

Several local environmental factors may also influence spatial differences in bleaching susceptibility. For example, local upwelling may cool warm surface waters, and high water flow resulting from currents and turbulence can result in reduced coral bleaching severity (West & Salm, 2003). Cloud cover (Mumby et al., 2001) and turbid water conditions can help decrease irradiance, but high sediment loads can cause smothering of coral polyps and reduced respiration, a decline in the photosynthetic productivity of zooxanthellae, and have several sublethal effects such as lowered calcification rates, damage to tissue, and reduced growth (Erftemeijer et al., 2012). Colony and reef-scale structural complexity may also work in sometimes opposing ways to influence the degree of thermal stress a reef experiences.

Highly rugose environments may be able to create shade within habitats, thereby helping to reduce temperatures in small pockets of the reef. However, colonies that have higher surface complexity and increased light-harvesting ability can also have higher susceptibility to bleaching as a result (Marcelino et al. 2013). In addition, colony microcomplexity can modify environmental conditions at small scales (Chamberlain and Graus 1975) thereby altering coral response to stress.

In addition to all these factors, the natural history and long-term environmental conditions of a particular reef system can also influence how coral taxa respond to disturbance, especially if historical conditions may have favored adaptation or tolerance to stress. Kāne‘ohe Bay, where this study was conducted, has experienced a unique history of human impacts (Hunter and Evans, 1995). From the 1940s to the 1970s, extensive dredging, land runoff, and the discharge of raw, untreated sewage in the southern section of the Bay led to anoxic and turbid conditions and the proliferation of *Dictyosphaeria cavernosa*, which outcompeted corals on many reefs (Evans 1995). These reefs were also subject to multiple freshwater kills in 1965, 1969, 1987 (Sukhraj 2014) and more recently, in 2014 (Bahr et al. 2015). After the diversion of the sewage outfall in 1977-78, however, coral cover began to increase, and stands today at nearly 60%. Their recovery is especially striking given that Hawaiian reefs have suffered from coral disease outbreaks in 2011, 2012, and 2015 (Jury & Toonen, 2019) and have been exposed to multiple coral bleaching events in 1996, 2002, 2004, and more recently, in 2014 and 2015 (Bahr et al., 2017). Additionally, the corals in Kāne‘ohe are living at temperatures 1-2 °C higher than surrounding waters and under elevated pCO_2_ conditions (Bahr et al., 2015), yet these parameters vary extensively across the Bay (Guadayol et al. 2014). This history of living under elevated stress is thought to be one of the main factors contributing to the resilience of these reefs (Jury and Toonen 2019). However, reef response appears to change with every bleaching event because of changes in local environmental condtions. For instance, Bahr et al. (2017) reported that patterns of bleaching and mortality in Kāne‘ohe Bay differed after 1996, 2014, and 2015, due to differences in the direction and magnitude of thermal stress and degree of freshwater inflow into the Bay prior to the bleaching event. Reef resilience is not static and understanding how changing conditions interact to affect patterns of coral bleaching and mortality – especially as temperature anomalies become more frequent – is a top conservation priority.

New tools and techniques like structure-from-motion photogrammetry (hereby, SfM) now enable robust and efficient quantification of reef structure and community change in ways that traditional techniques have not been able to accomplish before (Pizarro et al., 2017). Photogrammetry is the process of extracting quantitative information about a scene (in this case, a reef) through the collection of a series of 2-dimensional images (Burns et al. 2015; Pizarro et al., 2017). These overlapping 2D images can then be stitched together to estimate 3-dimensional structure and may be utilized to extract data on a range of metrics, from community demography, to several measures of surface complexity (rugosity, fractal dimension, and height range) and estimates of shelter space and volume (Edwards et al. 2017;Torres-Pulliza et al., 2020; Million et al., 2021). Because SfM generates high resolution orthophotos and photogrammetric products can be spatially registered, using this technique can be especially useful to track changes in colony health through time.

In this study, we tracked the fine-scale responses of 30 coral reefs throughout Kāne‘ohe Bay using SfM during a bleaching event in the summer and autumn months of 2019. Our primary goal was to quantify the spatial, temporal, and taxonomic variability in bleaching severity and mortality across the Bay. In addition, the spatial and temporal resolution of our data also allowed us to test hypotheses about the environmental and ecological factors that may have influenced these patterns. To assess this, we collected data on temperature, depth, sedimentation, and wave velocity, alongside estimates of habitat complexity. We hypothesized that differences in coral assemblage and reef-level environmental conditions would influence bleaching severity and mortality. In particular, based on previous work in Kāne‘ohe Bay, we expected bleaching to be lower in reefs that experienced conditions of lowered irradiance, potentially mediated by depth, the amount of suspended matter in the water column, and wave velocity (Bahr et al. 2017). We also expected bleaching severity and associated mortality to be highest in shallow reefs or those that experienced high temperatures, since the magnitude of thermal stress is a well known driver of the degree of bleaching and mortality (Eakin et al., 2010; Hughes et al., 2017). We predicted that reefs that were dominated by heat-susceptible species would experience more severe stress and elevated mortality (Loya et al., 2001) and expected the three-dimensional complexity of a reef to also influence the degree of bleaching and mortality it experienced.

## Methods

### Field methods

#### Data collection using Structure-from-motion photogrammetry (SfM)

We collected imagery at 30 patch reefs spread across the north-south extent of Kāne‘ohe Bay (Fig.1). These sites have been monitored since 2017 and represent a broad range of exposures and environmental conditions (Caruso et al., 2021). We started our surveys in August 2019 when we began to detect extensive coral paling due to rising temperatures.

**Figure. 1:**
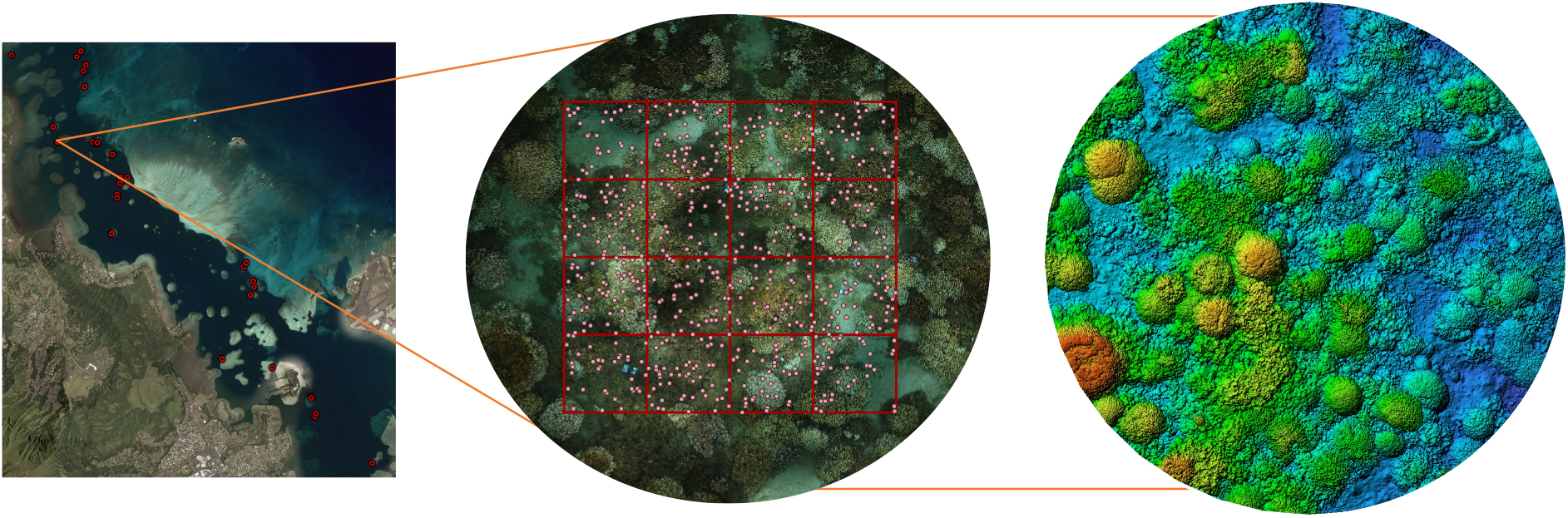
Schematic map of Kaneohe Bay showing 30 study sites and a corresponding orthomosaic and digital elevation model (DEM) for an individual site. 640 random points within an 8 × 8 m grid on the orthomosaic were annotated for substrate type at each site during the study period. Habitat complexity metrics were estimated from DEMs, where colours correspond to reef depth (red/green=shallow, blue=deep).

Our field methods have been described in detail in Roach et al. (2021). In short, we used a spool and line set-up to image approximately 112 m^2^, to create orthomosaics and digital elevation models (later cropped to 64 m^2^ for annotation). We used a Canon EOS Rebel SL3 DSLR in a waterproof housing with a wide angle 18-mm lens to ensure high overlap between images since these sites were all relatively shallow (2-5 m). Images were collected via SCUBA or snorkel depending on the depth of the site. The swimmer operating the camera system collected imagery by swimming in an outward spiral with the camera held roughly 1 m above the benthos, starting from the central point of the spool. Another individual held the center pole stationary over a cinder block used to mark the central point of these sites. Once the swimmer reached the outer edge of the spiral, they returned inwards in the same spiral swim pattern to the center, with the camera continuing to take pictures. An average of 3500-4500 images were collected per site; roughly 3 images/second. A site took approximately 20 minutes to map in the field. At each site, we placed 3 calibration markers in the area prior to imaging to help align images during processing. At each of these markers, we noted a) depth and b) compass bearing in relation to the center of the reef area being imaged. These were then used to accurately align and georeference models respectively.

We monitored each of the 30 patch reefs every 3-4 weeks between August 2019 -November 2019, which corresponded to peak bleaching period. Following this, we resurveyed sites in January 2020 and again in September 2020, one year after the bleaching event. This resulted in a total of 210 reef models across all sites and time points. We discarded 30 of these models due to holes in the model or poor-quality imagery.

#### Quantifying the environmental characteristics of reefs

Data on temperature, sediment, and wave energy were collected as part of a study on coral clonality and we refer the reader to this for further details (see Caruso et al., 2021). In short, temperature data was collected alongside photogrammetry imagery at each site at 10-minute intervals on Hobo Pendent or Water Temp Pro V2 loggers from the center of each site. We summarized hourly temperatures per day to calculate daily means. We used this to estimate Degree Heating Weeks (DHW) as time spent above the maximum monthly mean (MMM+1 degree, or 28° C for Kāne‘ohe Bay; Jury & Toonen, 2019). DHW have become a common predictor of coral bleaching, with significant bleaching usually occurring at over 4 DHW, and we use mean DHW in our analyses of bleaching and mortality. We used average temperature values from October 16^th^ -31^st^ for the entire month due to a gap in sampling in the first two weeks of October.

16” capped PVC pipes were used to collect sediment data every 1-2 months for each site between 2017-2019. Given that these sites differ significantly in their water residence times, these sedimentation rates are likely more indicative of the suspended particulate matter in the water column rather than how much new material is being deposited on a reef. Sediment data were not collected during the bleaching period, but we use mean sediment values for every site over the two-year duration of collection in our analyses.

Current meters were deployed at a cement block at a single site in every region in 2019 and used to calculate root mean square (RMS) wave velocity (Caruso et al., 2021). Regions were primarily defined based on water flow conditions, with sites in the north-northwest (region 5) having lowest water residence times (∼1 day), those in the central section of the Bay (regions 2, 3, 4) experiencing moderate water residence times of 2-10 days based on their relative exposures, while sites in the southernmost section (region 1) had significantly higher water residence times of 1-2 months (Lowe et al., 2009). These values were extrapolated to all sites within a region. Depth was measured from the center of each site.

### Data processing and analysis

#### Processing 3D reef models and measuring bleaching severity

Images for every site were aligned in *Agisoft Metashape Pro (Version 1*.*7*.*6)*. Prior to alignment, photographs that were out of focus or in blue water were manually removed. 2D orthophotos and gridded digital elevation models (DEMs) were created for each site following a standard workflow that begins with photo alignment, and leads to the creation of a 3D dense point cloud, from which textured meshes, DEMs, and orthophotos (also referred to as orthophoto mosaics, or orthomosaics) were outputted (Burns et al. 2015). Depth and compass measurements were used to accurately calibrate and georeference individual models. Completed orthomosaics were imported into *QGIS 3*.*16* for further analysis. All models were cropped to an 8 m x 8 m square, so that the area analyzed in every mosaic was 64 m^2^. 640 random points (or 10 per m^2^) were overlaid on the mosaic shapefile. For each point, we assessed benthic status in 22 categories (e.g., coral, cca, turf, macroalgae, rubble, dead skeleton; see Supplementary Table 1). When the substrate was a coral species, we assessed its bleaching status using the categories 0-3, where 0=healthy, 1=pale, 2=significantly bleached, 3=stark white. When a coral died, its identity changed most commonly to “dead skeleton”, “turf”, or “cca”. While we identified a total of nine coral species in our reef sites, *Porites compressa* and *Montipora capitata* make up the large majority (>90%) of the coral cover on many reefs here. We therefore restricted our analyses to individuals of these species and *Pocillopora*, that were abundant in a number of our survey sites. *Pocillopora* (*P*.*meandrina* and *P*.*damicornis*) were grouped by genus due to their diverse colony morphologies in Kāne‘ohe Bay and recent taxonomic reclassifications that make them difficult to accurately identify visually (Johnston et al., 2018). We tracked the same 640 points through time for every reef site (Fig.1). From this we summarized data on the community composition at every site and used a multivariate analysis to assess how the relative proportion of different substrates changed through the bleaching event.

DEMs were also exported into *QGIS* and cropped to the same size as orthomosaics (Fig.1). We divided them into 16 contiguous squares (covering an area of 64 m^2^) from which surface rugosity (R) and fractal dimension (D) were calculated for 2 × 2 m grids for every DEM, making up a total of 16 values per site, using the same methodology as Torres-Pulliza et al. (2021). R values were log-transformed, and all 16 R and D values from the first survey time were used in models assessing bleaching severity and mortality. Completed annotation shapefiles and habitat complexity metrics were imported into the statistical software R for further analysis (R Core Team, 2021).

#### Data analysis

We quantified the drivers of bleaching severity using a cumulative link mixed model since our response variable was categorial. The response variable in our model was bleaching severity (i.e., status 0-3). Our fixed effect variables were maximum DHW, depth, mean sediment, wave velocity, coral species, rugosity, and fractal dimension. Site was included as a random effect to account for site-level differences not explained by fixed effects. For model selection we built our full model, an environmental model with only environmental data (DHW, depth, sediment, wave velocity), and an ecological model with only our ecoloical data (rugosity, fractal dimension, species composition). We dropped non-significant fixed-effects terms sequentially, based on log-likelihood ratio tests. We selected our best model based on the lowest Akaike Information Criteria (AIC) values (Supplementary Table 4).

Models were fitted using “clmm” in the R package “ordinal” (Christenson 2018).

To identify the strongest predictors of mortality we used a generalized mixed effects model (glmer) with a binomial distribution. Our fixed effects were mean DHW, depth, mean sedimentation, wave velocity, fractal dimension, rugosity, and substrate, while site was a random effect. We used mean DHW as opposed to maximum DHW in this analysis to better capture change in temperature through time, but both mean and max had the same effect in our model. Similar to the clmm, we built our full model, an environmental model, and an ecological model separately. We dropped non-significant variables sequentially to get our best model (Supplementary Table 5).

## Results

### Coral bleaching and mortality

We recorded extensive coral bleaching during August-December 2019, with 80% of all corals showing some signs of bleaching in this period (Figs.2b,3a). Sea water temperatures in Kāne‘ohe Bay peaked in August and September 2019 (Fig.2a), but degree heating weeks (DHW) were highest in October (7° C week). Mean DHW between August-November was 5.3 (° C week). Sites varied from each other in their mean daily temperatures, which ranged from 26-31°C through the bleaching period (Fig.2a). Although many corals began to bleach in August when seawater temperatures were at their highest, peak bleaching coincided with maximum degree heating weeks in October (7 DHW, Fig.3a). Bleaching severity varied markedly with species, with *Pocillopora* the most severely affected, followed by *Porites compressa* and *Montipora capitata* (Fig.2b). *Pocillopora* generally bleached earliest and experienced a second peak in bleaching in some sites in November. *P*.*compressa* and *M*.*capitata* experienced similar patterns in bleaching severity, with the majority of surveyed corals in status 1 or pale (Fig. 2b).

**Figure. 2:**
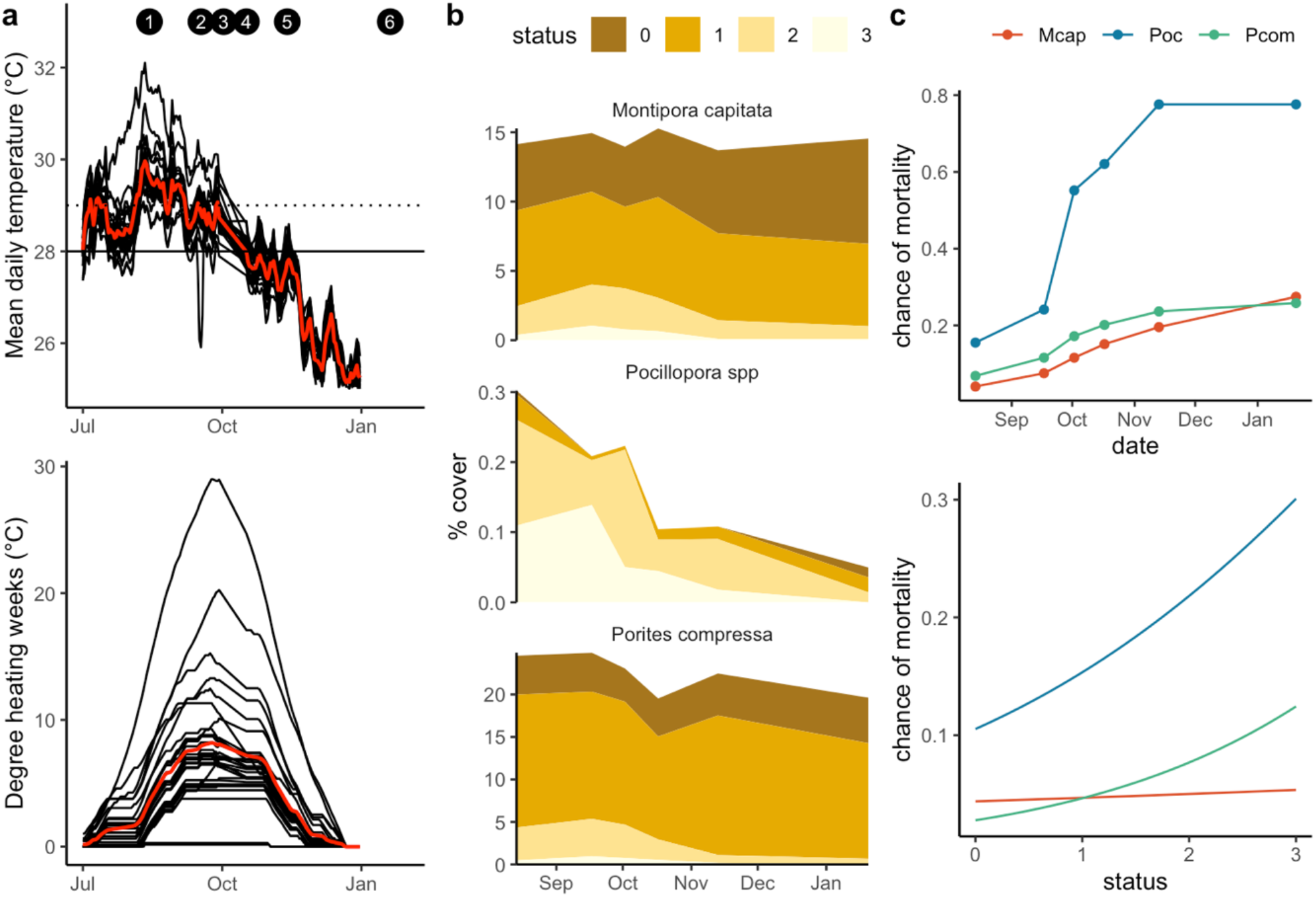
(a)Temperature profiles and DHW for all 30 sites, July 2019-January 2020. Red line indicates mean temperatures/mean DHW. Solid line corresponds to mean monthly maximum (MMM), dotted line indicates bleaching threshold (MMM+1 deg.C) (b) bleaching severity for 3 species, showing changes in percent cover over time. Status codes correspond to Fig.3. and (c) mortality (0=alive, 1=dead) for the three coral species over time and with increasing bleaching status (0=healthy, 3=stark white).

All three species we tracked experienced some mortality from bleaching. Mortality was lowest for *Porites compressa* (19%) followed by *Montipora capitata* (23%) and *Pocillopora* (77.5%) (Fig.2c). These rates are more indicative of the death of individual points on coral colonies rather than whole colony mortality. Mortality rates in this study are therefore best interpreted as a decrease in overall coral cover rather than the loss of entire colonies. In general, mortality peaked in October but continued to accumulate throughout the bleaching period (Fig.2c). However, maximum mortality for *M*.*capitata* occurred in January after temperatures had dropped. Mortality was strongly linked to bleaching stress, with the chance of mortality for all species increasing with bleaching status, peaking at status 3 (stark white).

### Compositional and environmental drivers

Bleaching severity and mortality were each influenced by species composition and local abiotic conditions, creating variation in the impacts of thermal stress across the study site (Figs.3, 4). Sedimentation rates varied substantially across reefs, ranging nearly 300-fold from 0.01 g/day to 2.93 g/day across sites (see Caruso et al. 2021). RMS (wave velocity) ranged from 0.8 to 12.56 cm/s. Sites in the northern sections of the Bay (Region 5) had higher sediment levels as well as higher RMS than sites in other regions (Fig. 3b). In contrast, sites in Region 1 clustered together based on their rugosity (R). Depth varied between 0.5-3.5 m across all reef sites (with a ∼0.4 m daily tidal fluctuation). In addition, sites differed in their relative cover of different substrates with live coral cover highest in Regions 2 and 4 (Fig 3c). Turf, bare coral skeleton, and sand and rubble made up over half of the cover in sites in Regions 1 and 3. Crustose coralline algae (CCA) had the highest cover in region 5 (Fig. 3c).

**Figure 3:**
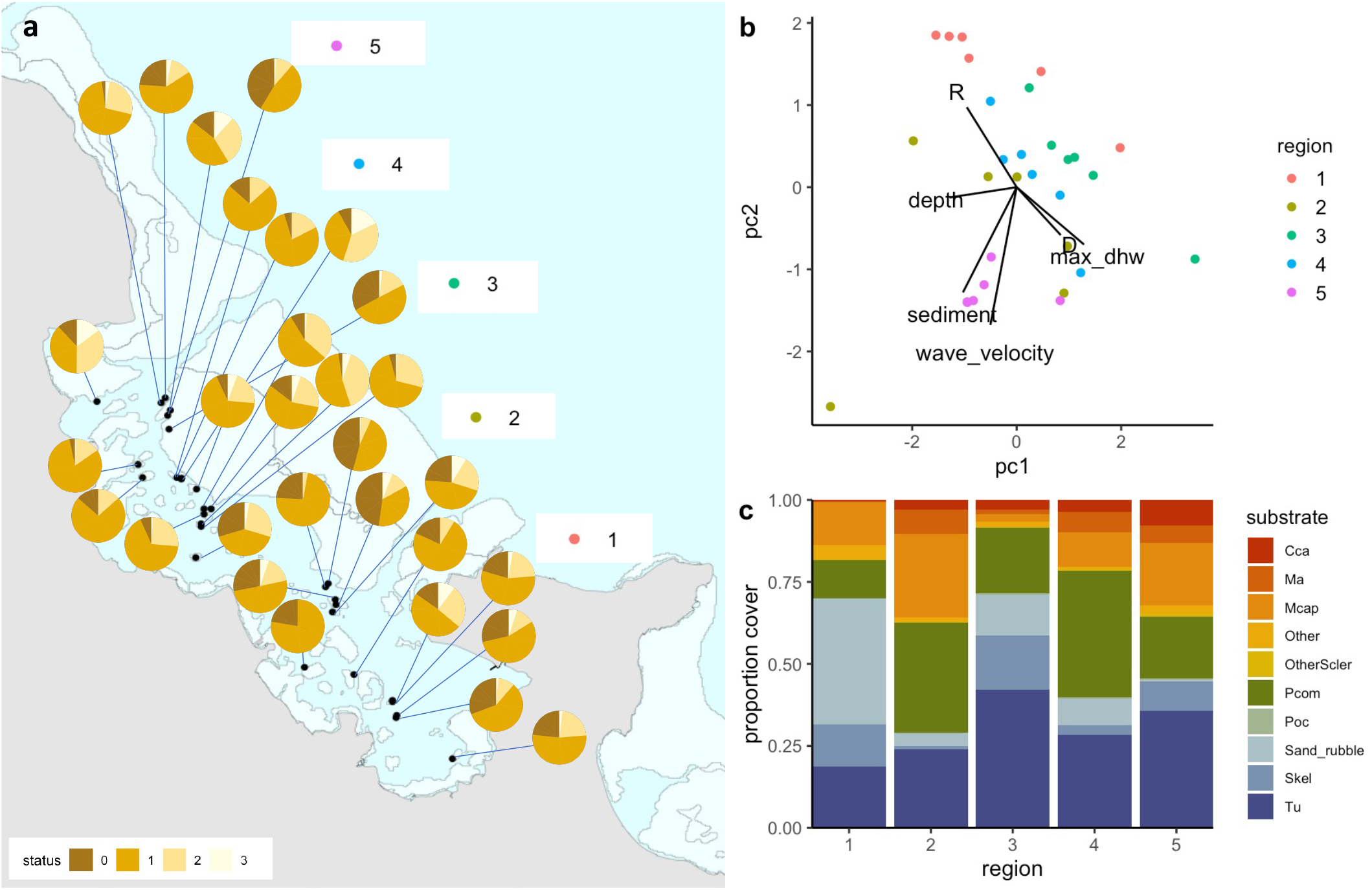
(a) pie charts showing proportion bleaching at all sites across 5 regions during peak bleaching (status corresponds to bleaching severity, where 0=healthy, 1=pale, 2=significant loss of pigmentation, 3=stark white), (b) PCA summarizing all environmental and ecological parameters measured, colored by region (R=rugosity, D= fractal dimension, max_dhw=maximum degree heating weeks), and (c) relative proportion of the dominant substrate types across the five regions in the Bay (cca=crustose coralline algae, ma=macroalgae, Mcap=Montipora capitata, Pcom=Porites compressa, Poc=Pocillopora, Skel=skeleton, Tu=turf).

**Figure 4:**
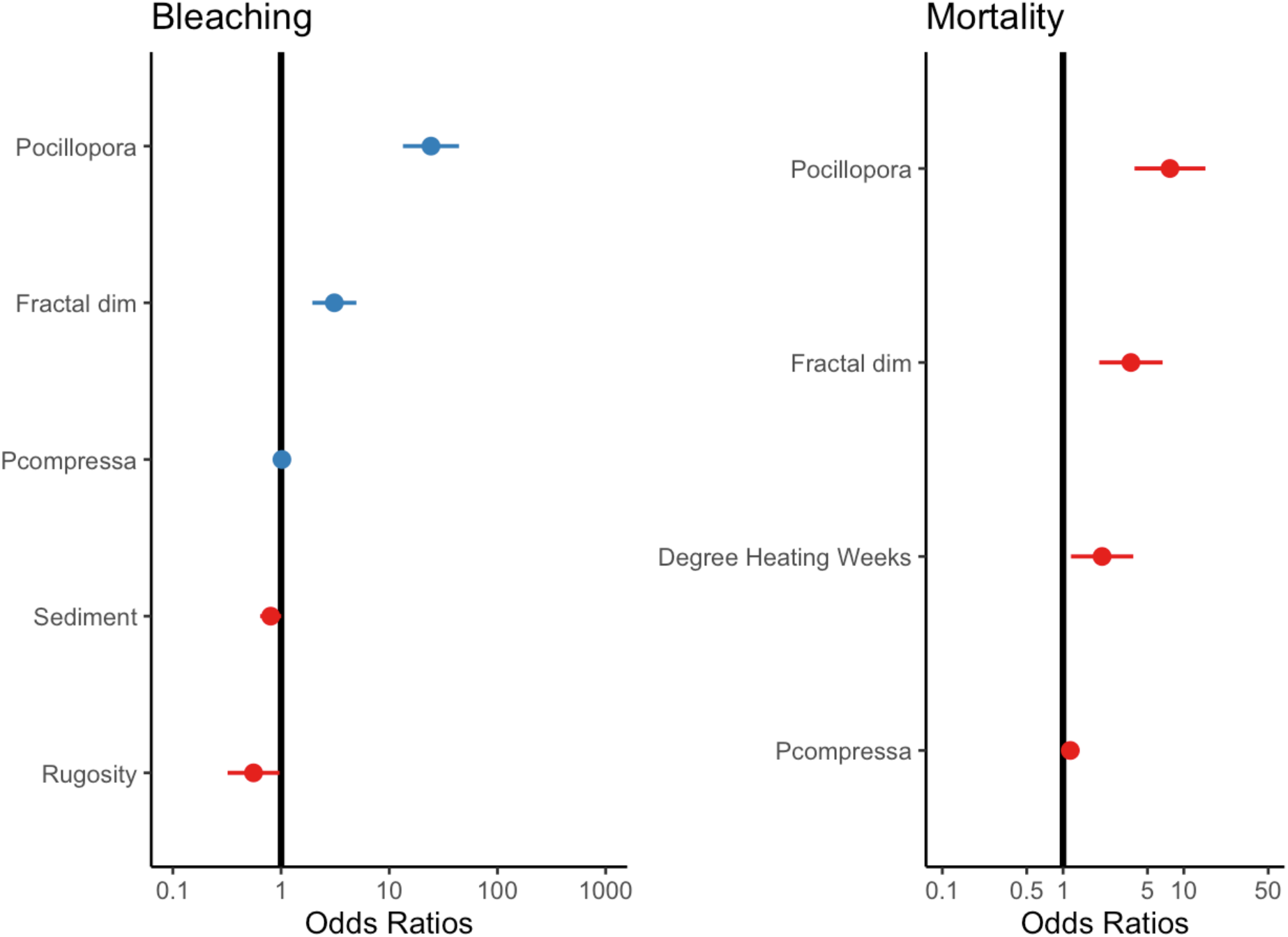
The effects of significant environmental and ecological variables on bleaching and mortality. Null line indicates no effect. Lines from individual points indicate confidence intervals. See Supplementary Fig. 4 for the full model.

Our analysis showed that habitat complexity (rugosity R, fractal dimension D) and coral species were important predictors of bleaching severity (Table 1). Coral assemblage had the largest effect on bleaching severity, with sites dominated by heat-sensitive *Pocillopora* experiencing more severe bleaching than others (p <0.001). Following this, sites that bleached more severely were associated with higher scores of fractal dimension D (p <0.001, Table 1). In contrast, there was a negative relationship between bleaching severity and rugosity (p <0.05), with more severe bleaching occurring in reefs with lower rugosity scores (Fig.4). Sediment loads had a slight negative effect on bleaching severity, with corals bleaching more severely in reefs with lower levels of sedimentation. We did not find an effect of depth, wave velocity, or DHW on bleaching severity (Supplementary Table 2).

**Table 1:** Parameters and coefficient estimates of fixed effects in the final cumulative link mixed model (clmm) with bleaching status as a categorical response variable. Full model in Supplementary Table 2.

| Variable | Estimate | SE | z Value | P Value |
| --- | --- | --- | --- | --- |
| Sediment load | -0.22102 | 0.11395 | -1.940 | 0.0524 . |
| Rugosity | -0.58611 | 0.28187 | -2.079 | 0.0376 * |
| Fractal dim | 1.13326 | 0.23916 | 4.738 | 2.15e-06 *** |
| *Pocillopora* spp | 3.18983 | 0.30445 | 10.477 | < 2e-16 *** |
| *Porites compressa* | 0.01797 | 0.06249 | 0.288 | 0.7737 |

Coral mortality was strongly affected by thermal stress, coral species composition, and fractal dimension. Predictably, heat-susceptible *Pocillopora* suffered maximum mortality during this event and had the largest effect in our model (Table 2). Notably, fractal dimension (D) had the same effect on mortality as it did on bleaching severity, with sites with higher D experiencing more mortality. Variability in average thermal stress across reefs also impacted mortality rates, with mortality increasing in reefs that experienced a higher number of degree heating weeks (Fig. 4).

**Table 2:** Summary statistics of the best fit glmer predicting the probability of coral mortality. Fixed effects were degree heating weeks (DHW), substrate (Pocillopora spp, Montipora capitata, Porites compressa) and Fractal dimension (fractal dim). Full model in Supplementary Table 3.

| Variables | Estimate | SE | z Value | P Value |
| --- | --- | --- | --- | --- |
| Intercept | -3.95635 | 0.81689 | -4.843 | 1.28e-06 *** |
| DHW | 0.74366 | 0.30362 | 2.449 | 0.0143 * |
| *Pocillopora* spp | 2.03826 | 0.34506 | 5.907 | 3.48e-09 *** |
| *Porites compressa* | 0.13781 | 0.07624 | 1.807 | 0.0707 . |
| Fractal dim | 1.29318 | 0.30840 | 4.193 | 2.75e-05 *** |

The 3D mosaics resulting from our imagery allowed us to quantify fine-scale compositional changes across a large area (30 sites, and 19200 points covering 1920 sq m of area). By quantifying proportional change in the cover of key taxonomic groups at each site, we identified a marked shift in the composition of reefs as mortality progressed (Fig.5a,b). Moreover, even though sites varied considerably in their initial compositions, there was a common trend towards an increase in benthic algal cover post-mortality (Fig.5c). A decrease in coral cover was accompanied by an increase in the cover of turf and macroalgae (Fig.5b). We did not note any significant changes in the three-dimensional structural complexity of reefs during this one year period (Supplementary Fig.5).

**Figure 5:**
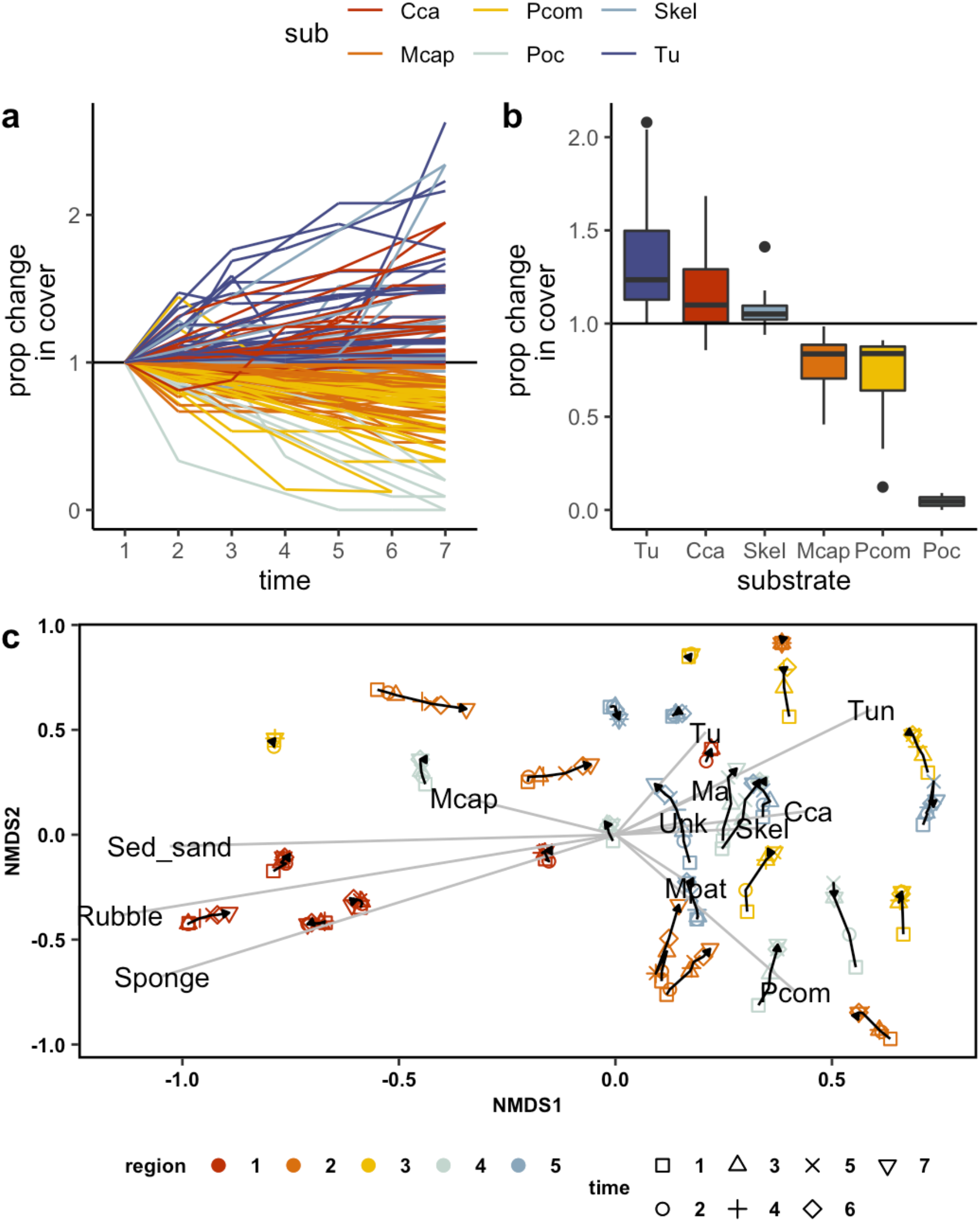
Changes in the proportional cover of different substrates over the course of the bleaching event. (a) an increase in turf (Tu) and CCA was associated with a decrease in P.compressa (Pcom) and Pocillopora (Pdam) during October-November (times 3, 4, 5) (b) changes in % cover by substrate type over the bleaching event, (c) NMDS showing a shift towards Tu (turf), Ma (macroalgae), Skel (skeleton), and CCA through survey times across the study regions.

## Discussion

The results of this study show that corals in Kāne‘ohe Bay experienced a moderate to severe bleaching event in 2019, with up to 80% of corals showing some signs of stress during this period. In general, reefs around O’ahu did bleach more extensively (∼43%) than other Hawaiian islands during the 2019 event (Winston et al. 2020). Variation in bleahing severity in Kāne‘ohe Bay was mediated by sediment levels and species composition, while mortality was driven largely by temperature and the identity of the coral.

### Severity and extent of bleaching

Peak bleaching stress in all parts of the Bay occurred in the month of October – while temperatures were highest in August and September, October had the highest degree heating weeks (7 DHW). This follows a similar pattern to the bleaching events in 2014 and 2015, when DHW were highest in September and October (Ritson-Williams & Gates, 2020).

However, cumulative thermal stress over the bleaching period was lower in 2019 (this study) at 5.3 DHW over the 4-month bleaching period, as opposed to 6 DHW in 2014 and 12 DHW in 2015. Corals generally begin to show significant signs of stress at above 4 DHW; however, in 2019, many corals began to bleach in August when DHW were only 3.3. After rising to 32°C in August, seawater temperatures dropped to 27°C in November, and to 25.5°C by December. In general, Kāne‘ohe bay experiences significant fluctuations in SSTs, swinging from 19°C in winter months to 29.3°C during normal summers (Caruso et al., 2021; Ritson-Williams & Gates, 2020). In the past, temperatures have crossed 30°C only in 1996, 2014, and 2015, for a maximum of 17 days in 2015. During this bleaching event, some sites experienced temperatures above 30°C for an entire month.

The overall rates of bleaching we record in this study are higher than those recorded in 2014 (∼65%) and 2015 (∼50%) (Bahr et al., 2017). However, it is worth noting that many corals in Kāne‘ohe Bay appear pale through much of the year, and our cumulative bleaching estimates likely reflect that. Previously, surveys have often been conducted following peak temperatures and not at the granularity of the present study, which could have resulted in lower estimates of past bleaching. Notably, though, average mortality for *P*.*compressa* and *M*.*capitata* (21%) was slightly lower than cumulative mortality following 2015 (22%).

Increased rates of bleaching but lower average mortality suggests some degree of acclimatization to repeated thermal stress in these species. Bleaching in 2019 followed a similar pattern to 2015 with sites in the northern region bleaching most severely followed by those in the central and southern sections of the Bay (Bahr et al., 2017).

Bleaching was variable for species. *Pocillopora* bleached most severely in the Bay, followed by *P*.*compressa* and *M*.*capitata. Pocillopora* has been severely affected by past bleaching events, and these patterns continue with our study (Bahr et al., 2017). While a slightly larger proportion of *P*.compressa bleached than *M*.*capitata*, the latter had a higher proportion that bleached more severely. However, *M*.*capitata* that recovered seemed to do so faster than *P*.*compressa*. This is in contrast to patterns following the bleaching event of 2015, where *P*.*compressa* visually recovered more rapidly than *M*.*capitata* (Matsuda et al. 2021). Higher than normal nutrient and zooplankton concentrations associated with elevated levels of sedimentation in Kāne‘ohe Bay could assist heterotrophically plastic corals like *M*.*capitata* to meet their metabolic demands and survive through periods of elevated sea water temperatures. This could be one of the reasons this species was able to recover pigmentation faster than *P*.*compressa* after this bleaching event. However, *P*.*compressa* experienced less mortality than *M*.*capitata*, even if it showed a slightly longer lag in recovery time. In contrast, while *Pocillopora* showed greatest mortality during this event, colonies that did not die recovered faster than both *P*.*compressa* or *M*.*capitata*.

### Environmental drivers of bleaching

Several studies have confirmed that anomalously high temperatures are the leading cause of coral mortality and habitat loss worldwide (Sheppard, 2003; Donner et al., 2005). However, local reef conditions can work to mediate the effects of elevated temperature, thereby influencing the response of corals to thermal stress. We find that site-level differences in habitat complexity and assemblage type played an important role in determining bleaching severity during 2019, with reefs that were dominated by *Pocillopora* or those that had lower sediment loads experiencing more severe bleaching than others (Fig.4).

The structural complexity of a reef and its importance for biodiversity has been recognized for decades, but how underlying reef structure can reinforce or buffer the effects of bleaching and mortality is less known (Ferrari et al., 2016). Our study finds that higher levels of complexity at the colony-level scale (i.e., rugosity) were associated with reduced bleaching severity. This pattern might be attributed to the prevalence of heat-resistant *Porites compressa* across these sites, which experienced the least bleaching during this event. In addition, *P*.*compressa* grow in large, mounding colonies in the Bay often forming monospecific stands and contributing to much of the larger-scale 3D complexity on these patch reefs (Supp Fig1a). Reef patches with higher rugosities might also have been able to create more shade within the reef, which could have further protected colonies or parts of colonies from thermal stress. In contrast, higher scores of fractal dimension – which are more indicative of complexity at the micro-scale – were related to higher levels of bleaching severity, potentially reflecting the effect of bleaching on heat-susceptible *Pocillopora* spp.

*Pocillopora spp* are intricately branched and were associated with higher D scores than *P*.*compressa* and *M*.*capitata* (Supp Fig1b). Fractal dimension and rugosity capture different elements of complexity on a reef and might work in asynchronous ways to affect processes like bleaching and mortality (Torres-Pulliza et al., 2020). While coral diversity in Kāne‘ohe Bay is low, species exist in a range of morphotypes and contribute differently to overall reef structure (Miller et al., 2021). While we did not detect changes in the 3D complexity of reefs one year post-bleaching, further monitoring will provide insight into how reef-scale R and D change as corals of different species grow, recover, or die.

Sedimentation has long been known to be a major driver of reef health in Kāne‘ohe Bay due to its history of extensive dredging, coastal development, and land run off from adjoining coastal areas. Variation in sediment loads across sites played a moderate role in determining bleaching response. While higher sediment loads have generally been found to be damaging to reef corals (Erftemeijer et al., 2012), studies have also found that the resulting turbidity may act to shield corals from radiation and the effects of increased temperature (Anthony et al., 2007). The tolerance of different coral species to sediment may vary based on their growth forms, with some morphologies -like hemispherical or columnar colonies -more efficient passive shedders than others (Riegl 1996). On the other hand, some colony shapes may be able to create vortices that help to flush sediment out of colonies in areas of high water flow (Riegl et al., 1996). Following the bleaching event of 1996, areas of Kāne‘ohe Bay near stream mouths suffered little or no bleaching, even though corals experienced the same temperatures as in other parts of the Bay (Jokiel et al. 2004). In our study, elevated levels of sediment reduced bleaching stress potentially by reducing damage from UV radiation. Increased sediment loads in the water column could also be indicative of a higher degree of mixing and movement of water, which might have helped reduce temperatures at local reef scales. However, wave velocity did not have a significant effect on bleaching severity.

### Mortality: causes and consequences

All three coral species experienced mortality during this bleaching event. Species mortality in this study (77.5% for *Pocillopora*, 19% for *P*.*compressa*, 23% for *M*.*capitata*) were on par with the 2015 bleaching event. However, our estimates of mortality also include partial colony mortality, especially for *P*.*compressa* and *M*.*capitata* due to the large size of colonies in Kāne‘ohe Bay. Nevertheless, *Pocillopora* generally suffered whole-colony mortality.

Higher overall rates of mortality indicate that the extent and duration of the 2019 bleaching event was more severe for these species than expected. While mortality did not lead to changes in the three-dimensional complexity of these reefs in the study period, we observed a rapid takeover of dead corals by algae (turf, cca). This trend towards algae was consistent across reefs, even though they differed in their coral community composition.

Species differences and variation in fractal dimension D between reefs largely drove mortality rates, alongside some fine-scale spatial differences in thermal stress (Fig.4). While high SSTs and degree heating weeks are known to be major drivers of bleaching and mortality, it is striking that differences in DHW existed at the spatial scale of this study.

Average DHWs ranged from less than 1 to 24 through the bleaching period across sites, with sites in the northern region of the Bay (5) generally experiencing higher DHW (∼10) than those in the south and central regions (∼5 DHW). Sites that were more inshore such as those in region 3 also experienced higher DHW (∼9), potentially as a result of their higher water retention times. Lowest DHW occured in sites that were close to a channel and slightly offshore. Highest mortality rates coincided with sites that experienced greater than 8 DHW (Supp Fig.2). Similar to its effect on bleaching severity, higher fractal dimension was associated with higher mortality. While higher sediment levels and RMS were associated with decreased mortality rates, they did not have a significant effect on it. In general, temperature, species thermal tolerance and fractal dimension overtook the effect of all other environmental variables in determining patterns of mortality (Supp Fig. 3).

*Pocillopora* are some of the most susceptible species in the Bay and were among the worst impacted in Kāne‘ohe following the bleaching events of 2014-15, when 80-100% of monitored colonies bleached, and 19% had died by 2016 (Ritson-Williams & Gates, 2020). Branching coral taxa like *Acropora* and *Pocillopora* are generally considered to be more bleaching susceptible than massive forms such as *Porites* and tank experiments on *P*.*acuta* collected in Kāne‘ohe Bay have been found them to be susceptible to elevated temperature conditions (Bahr et al., 2020). Declines in *Pocillopora* raise questions about the larger functional consequences of altered coral assemblages in Kāne‘ohe Bay in an era of climate change. Early reports from the Bay documented over 20 common coral species here (Maragos 1972); however, only 14 of these species were observed on a resurvey of the same sites in 2009 (Sukhraj 2014, pp.124). While there have been taxonomic reclassifications of some of these species over time, genera such as *Porites* and *Montipora* appear to have been able to better acclimate or adapt to the unique conditions of this Bay and its historic disturbances, while others have likely declined in this period. While *Pocillopora*-like species are generally fast growing, have early maturity (Baird and Maynard 2008), and have been able to adapt to changing conditions in other places (Guest et al., 2016) their abundance, distribution, and rates of recruitment in Kāne‘ohe Bay will need further monitoring. Local conditions affect taxa differently, and it is conceivable that *Pocillopora* will see further declines in these habitats in the future or remain restricted to deeper reefs or reef patches where conditions are more favorable.

## Conclusions

Our study finds significant spatial heterogeneity in patterns of bleaching and mortality, that were driven by differences in environment (temperature, sediment), and ecology (reef complexity, coral assemblage). Our study provides a fine-scale assessment of bleaching and mortality by using methods that allowed us to conduct repeat surveys of the same reefs as they altered through time. The use of SfM in this study has helped create a permanent digital baseline of these patch reefs for future work. Continuing to track reef 3D reef metrics at different scales will provide greater insight into the mechanisms that help maintain structural complexity in the face of future warming. While the majority of bleached corals were able to recover after this event, *Pocillopora* spp. suffered significant mortality. Kāne‘ohe Bay has long been considered an ecosystem under stress where some coral species are still able to adapt and thrive, but future work will need to elucidate how heat-susceptible taxa contribute to the functional diversity of corals in the Bay and what the consequences of their decline will be on associated reef fish species and reef function.

## Acknowledgements

We would like to thank many researchers and volunteers who were critical to the success of this work – specifically, Kira Hughes, Devynn Wulstein, Annie Innes-Gold, Mollie Asbury, Damarias Torres-Pulliza, Valerie Kahkejian, Elizabeth Madin, and Shayle Matsuda. This work was supported by a NOAA grant (award number NOAA-NFA-NFAPO-2018-2005418).

