## Supplementary Information for "Fine-scale variability in coral bleaching and mortality during a marine heatwave"

Table 1: Description of substrate types assessed during coral annotation. Note that Pdam and Pmea were collapsed into Pocillopora spp. in our analyses.

| **Code** | **Description** |
| --- | --- |
| Sed_sand | Sediment and sand |
| Rubble | Rubble |
| Plob | Porites lobata |
| Mcap | Montipora capitata |
| Pdam | Pocillopora damicornis |
| Tu | Turf |
| Ema | Encrusting macroalgae |
| Brma | Brown macroalgae |
| Rdma | Red macroalgae |
| Pav | Pavona spp. |
| Plut | Porites lutea |
| Pmea | Pocillopora meandrina |
| Sp | Sponge |
| Tun | Tunicate |
| Bi | Bivalve |
| Cca | Crustose coralline algae |
| Mfla | Montipora flabellata |
| Mpat | Montipora patula |
| Pcom | Porites compressa |
| Losc | Lobactis scutaria |
| Lpur | Leptastrea purpurea |
| Skel | Dead coral skeleton |
| Coce | Cyphastrea ocellina |
| Lpap | Leptoseris papyracea |
| Unknown | Unknown substrate |

Table 2: Full clmm predicting bleaching severity showing all variables tested

|  | **status** | | |
| --- | --- | --- | --- |
| *Predictors* | *Odds Ratios* | *CI* | *p* |
| 0\|1 | 3.59 | 0.46 – 27.77 | 0.221 |
| 1\|2 | 85.17 | 10.97 – 660.99 | **<0.001** |
| 2\|3 | 607.15 | 77.96 – 4728.18 | **<0.001** |
| Max_DHW | 0.98 | 0.93 – 1.03 | 0.364 |
| Depth | 1.02 | 0.75 – 1.40 | 0.889 |
| Sediment | 0.67 | 0.48 – 0.92 | **0.014** |
| RMS | 1.07 | 0.98 – 1.17 | 0.114 |
| sub [Pocillopora spp] | 24.23 | 13.35 – 44.00 | **<0.001** |
| sub [Porites compressa] | 1.02 | 0.90 – 1.15 | 0.782 |
| R | 0.56 | 0.32 – 0.98 | **0.041** |
| D | 3.11 | 1.95 – 4.98 | **<0.001** |
| **Random Effects** | | | |
| σ^2^ | 3.29 | | |
| τ_00_ _site_ | 0.52 | | |
| ICC | 0.14 | | |
| N _site_ | 30 | | |
| Observations | 7565 | | |
| Marginal R^2^ / Conditional R^2^ | 0.064 / 0.191 | | |

Table 3: Full glmer predicting mortality showing all variables tested

|  | **mort** | | |
| --- | --- | --- | --- |
| *Predictors* | *Odds Ratios* | *CI* | *p* |
| (Intercept) | 0.02 | 0.00 – 0.18 | **0.001** |
| depth | 1.22 | 0.67 – 2.22 | 0.519 |
| mean dhw | 2.25 | 1.23 – 4.11 | **0.009** |
| Sed Mean [log] | 1.07 | 0.53 – 2.16 | 0.854 |
| rms | 0.66 | 0.30 – 1.46 | 0.306 |
| D analysis | 3.61 | 1.95 – 6.67 | **<0.001** |
| R log10 analysis | 1.10 | 0.55 – 2.19 | 0.791 |
| sub [Pocillopora spp] | 7.72 | 3.92 – 15.18 | **<0.001** |
| sub [Porites compressa] | 1.15 | 0.99 – 1.34 | 0.067 |
| **Random Effects** | | | |
| σ^2^ | 3.29 | | |
| τ_00_ _site_ | 2.55 | | |
| ICC | 0.44 | | |
| N _site_ | 30 | | |
| Observations | 7565 | | |
| Marginal R^2^ / Conditional R^2^ | 0.120 / 0.504 | | |

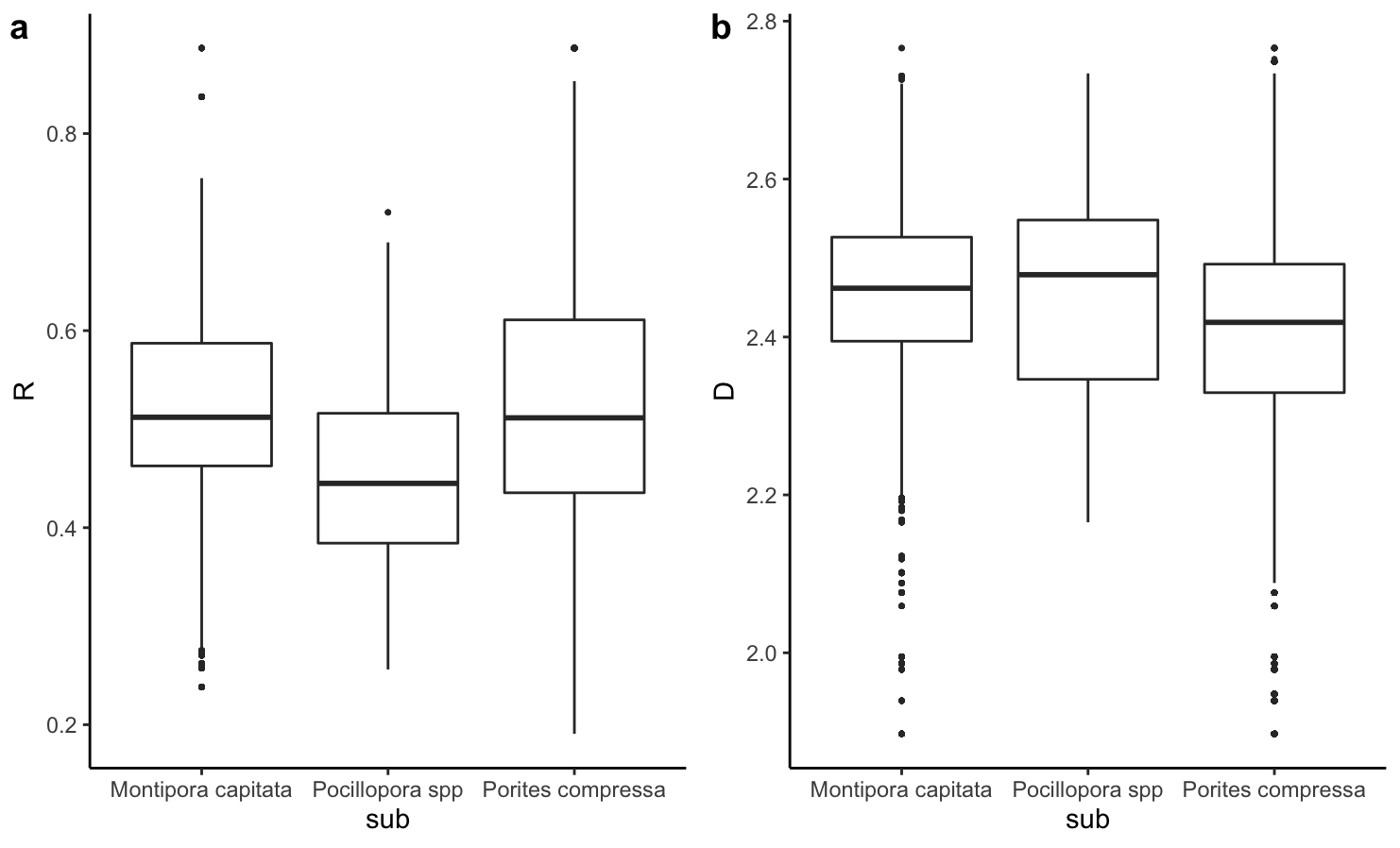

Fig.1: Relationship between coral taxon and a) rugosity R and b) fractal dimension D

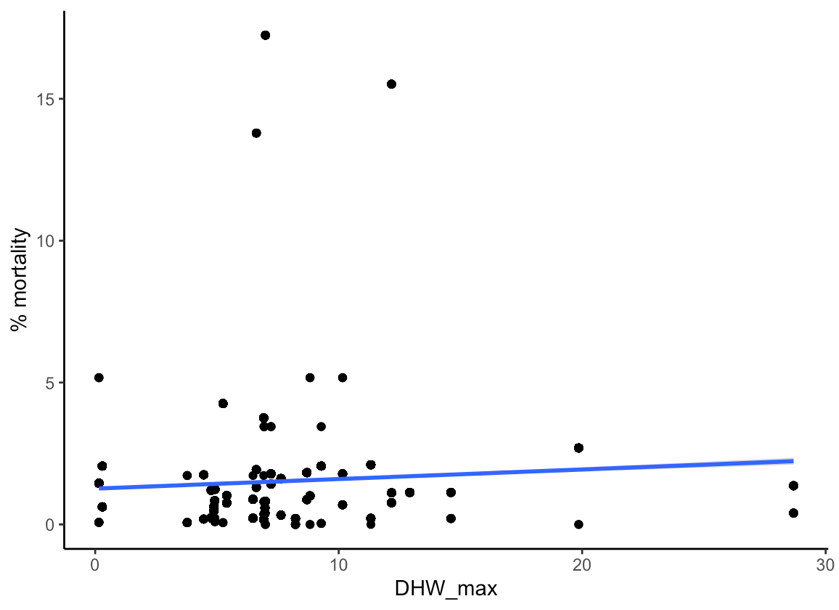

Fig.2: Max DHW vs % mortality across sites

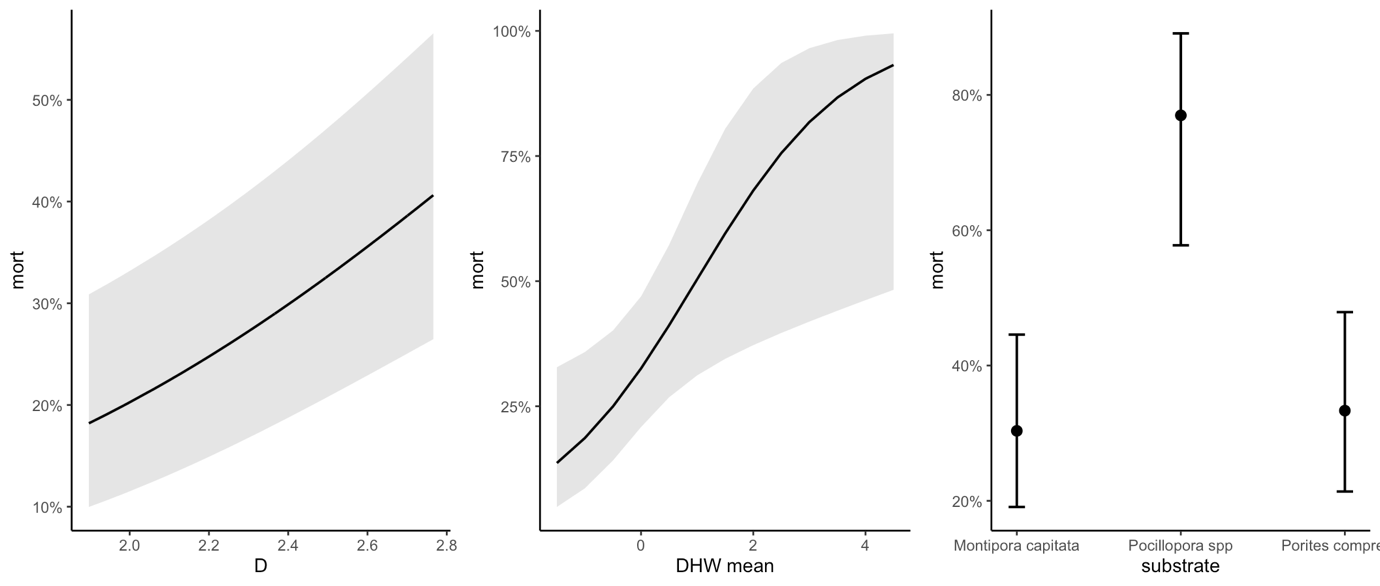

Fig.3: Modelled relationships between significant explanatory variables and percent mortality, showing predicted values for each model term.

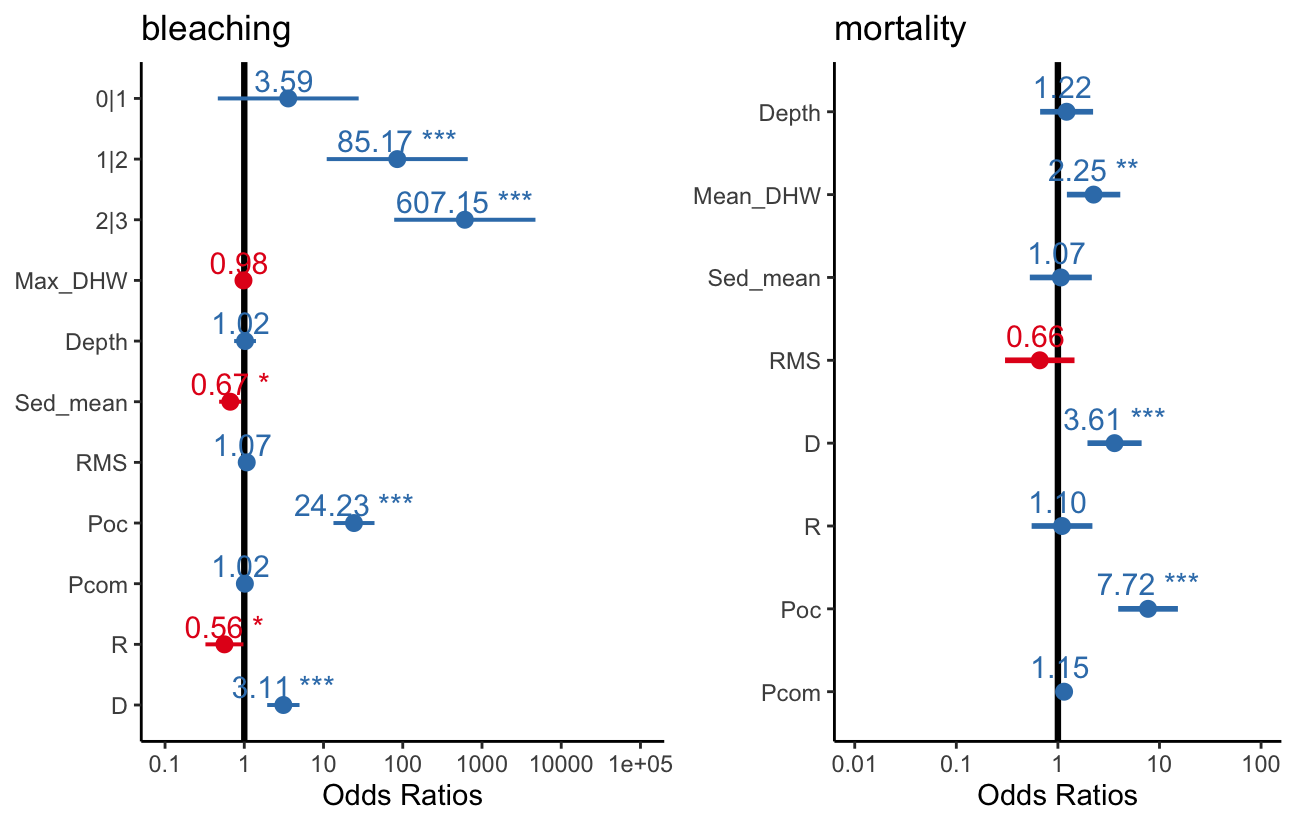

Fig 4: Effects plots for bleaching and mortality, showing the effects of all variables tested

Fig.5: Mean R and D over the course of the survey period

Table 4: AIC scores for models predicting bleaching severity

**Models	AIC**

Environmental model

Mean dhw + depth + sed + rms	15824

Ecological model:

Sub + R + D	15675

Best model:

Sed + R + D + sub	15672

Table 5: AIC scores for models predicting coral mortality

**Models	AIC**

Mean_dhw + depth + sed + rms	7634

Ecological model:

Sub + R+ D	7583

Best model:

Mean_dhw + D + sub	7577
